# Transcriptional Dynamics and Chromatin Accessibility in the Regulation of Shade-Responsive Genes in Arabidopsis

**DOI:** 10.1101/2025.01.23.634457

**Authors:** Sandi Paulisic, Alessandra Boccaccini, René Dreos, Giovanna Ambrosini, Nicolas Guex, Ruben Maximilian Benstei, Markus Schmid, Christian Fankhauser

**Affiliations:** Center for Integrative Genomics, Genopode building, University of Lausanne, 1015, Lausanne, Switzerland; Bioinformatics Competence Centre, EPFL, Lausanne, Switzerland; Bioinformatics Competence Centre, University of Lausanne, Lausanne, Switzerland; Umeå Plant Science Centre, Department of Plant Physiology, Umeå University, SE-901 87 Umeå, Sweden; Department of Plant Biology, Linnean Center for Plant Biology, Swedish University of Agricultural Sciences, S-750 07 Uppsala, Sweden; Unit of Food Science and Nutrition, Department of Science and Technology for Humans and the Environment, Università Campus Bio-Medico di Roma, 00128 Rome, Italy; HAYA Therapeutics SA, Route De La Corniche 5B, Batiment Alanine, 1066 Epalinges, Lausanne, Switzerland

**Keywords:** chromatin accessibility, transcription, shade, PIF7, Arabidopsis, ATAC-seq

## Abstract

Open chromatin regions host DNA regulatory motifs that are accessible to transcription factors and the transcriptional machinery. In Arabidopsis, responses to light are heavily regulated at the transcriptional level. Shade, for example, can limit photosynthesis and is rapidly perceived by phytochromes as a reduction of red to far-red light ratio (LRFR). Under shade, phytochromes become inactive, enabling PHYTOCHROME INTERACTING FACTORs (PIFs), particularly PIF7, to promote genome-wide reprogramming essential for LRFR responses. An initial strong and fast regulation of shade-responsive genes is followed by attenuation of this response under prolonged shade. We wanted to determine whether the transcriptional response to shade depends on chromatin accessibility. For this, we used ATAC-seq to profile the chromatin of seedlings exposed to short (1h) and long (25h) simulated shade. We found that PIF7 binding sites were accessible for most early target genes before LRFR treatment. The transcription pattern of most acute shade-responsive genes correlated with a rapid increase in PIF levels and chromatin association at 1h, and its decrease at 25h of shade exposure. For a small subset of acutely responding genes, PIFs also modulate chromatin accessibility at their binding sites early and/or late in the response to shade. Our results suggest that in seedlings a state of open chromatin conformation allows PIFs to easily access and recognize their binding motifs, rapidly initiating gene expression triggered by shade. This transcriptional response primarily depends on a transient increase in PIF stability and gene occupancy, accompanied by changes in chromatin accessibility in a minority of genes.

## INTRODUCTION

In Arabidopsis, a battery of photoreceptors is employed to sense the presence of neighbors by monitoring light quality (Ballare and Pierik, 2017, Fiorucci and Fankhauser, 2017, Pierik and Testerink, 2014). In dense plant communities, phytochromes perceive the neighbor threat as lower red (R) to far-red (FR) light ratio (LRFR), which signals the risk of being outgrown and shaded by neighbors. The pool of active phyB is then reduced and lifts the repression of PHYTOCHROME INTERACTING FACTORS (PIFs) to promote the shade avoidance response (Leivar and Quail, 2011, Chen and Chory, 2011). Active phyB inhibits PIF function by controlling their rapid turnover (e.g., PIF4 and PIF5), and DNA binding and/or transactivation activity (Shen et al., 2008, Shen et al., 2007, Park et al., 2012, Ni et al., 2013, Park et al., 2018, Leivar et al., 2020, Yoo et al., 2021, Xie et al., 2023). However, PIF7 regulation by phyB differs, as it does not involve rapid turnover. This may involve UBIQUITIN-SPECIFIC PROTEASE12 (UBP12) and UBP13 which promote de-ubiquitination of PIF7 (Zhou et al., 2021).

Transcriptional responses to neighbor threat (LRFR) are tightly regulated by PIFs, with a major role established for PIF7 (Li et al., 2012, Yang et al., 2023). Exposure to LRFR triggers accumulation and binding of PIFs to G-box and PBE-box motifs initiating rapid expression reprogramming. This affects numerous genes, including transcriptional regulators and auxin biosynthesis and signaling genes, essential for organ elongation and leaf repositioning (Devlin et al., 2003, Sessa et al., 2005, Bou-Torrent et al., 2008, Cifuentes-Esquivel et al., 2013, Leivar and Monte, 2014, Kohnen et al., 2016, Roig-Villanova et al., 2007, Hornitschek et al., 2012, Li et al., 2012, Tao et al., 2008). Transcriptomic changes induced by LRFR are very fast, reaching a peak between 15-90 minutes (Kohnen et al., 2016, Ciolfi et al., 2013) and are followed by relatively fast physiological adaptations (Fernandez-Milmanda and Ballare, 2021, Casal and Fankhauser, 2023, Gautrat et al., 2024, Krahmer and Fankhauser, 2024).

Upon perception of LRFR, PIF4, PIF5, and PIF7 induce expression of YUCCA genes to raise auxin levels (Won et al., 2011, Hornitschek et al., 2012, Li et al., 2012). Auxin produced in the cotyledons of seedlings is subsequently transported to the hypocotyl, where it promotes cell elongation (Procko et al., 2016, Procko et al., 2014). In an analogous process, auxin synthesized in the leaf blade (Zhao, 2012) is transported to the petiole, inducing elongation and leaf hyponasty (Iglesias et al., 2018, Kohnen et al., 2016, Michaud et al., 2017, Muller-Moule et al., 2016, Pantazopoulou et al., 2017). Within the cell, plants use polar auxin efflux carriers to establish polar auxin gradient, required for shade-induced hypocotyl, stem and petiole elongation, as well as leaf hyponasty (Kupers et al., 2023). Shade induced transcriptional responses and how they control organ growth and repositioning are quite well understood in Arabidopsis (Ballare, 2014, Ballare and Pierik, 2017, Buti et al., 2020, Ciolfi et al., 2013, Fiorucci and Fankhauser, 2017, Kohnen et al., 2016, Kupers et al., 2018). However, the epigenetic mechanisms and chromatin landscape underlying transcriptional regulation remain to be fully investigated.

The roles of H3K4me3 and H3K9ac in gene regulation have often been associated with actively transcribing genes. Among recent studies, Calderon et al have found that the H3K4me3 levels correlate with the active transcription of PIF-regulated and shade-induced genes (Calderon et al., 2022). Transcript levels of these genes increase before the H3K4me3 levels, implying that H3K4me3 increases as a consequence of active transcription, rather than preceding it, which is consistent with previous research, where high transcriptional activity of a particular gene locus was shown to induce the accumulation of H3K4me3 (Le Martelot et al., 2012, Kuang et al., 2014, Calderon et al., 2022). Shade also induces a significant increase in H3K9ac levels depending on PIF4, PIF5 and/or PIF7 (absent in the triple *pif4pif5pif7* or *pif457* mutant) (Willige et al., 2021). H3K9 hyperacetylation was observed not only on the gene bodies of the PIF7 targets, but also in the regulatory regions of specific genes such as *ATHB2* (Boycheva et al., 2024, Willige et al., 2021, Barnes et al., 2019). Interestingly, H3K9 hyperacetylation at PIF7 regulatory regions and target genes precedes H3K4me3 accumulation (Calderon et al., 2022, Willige et al., 2021).

Besides histone modifications, the chromatin landscape is heavily modified by ATP-dependent SWI/SNF-type chromatin remodeling complexes (Bieluszewski et al., 2023). These complexes adjust the position and occupancy of nucleosomes through sliding, eviction, or nucleosome deposition, thereby controlling the accessibility of DNA regulatory regions and, ultimately, gene transcription (Clapier et al., 2017). For instance, the SWI/SNF-type complex characterized by INO80 replaces the histone variant H2A.Z with the canonical H2A, in contrast to SWR1-containing complexes, which mediate H2A.Z deposition (Papamichos-Chronakis et al., 2011, Alatwi and Downs, 2015, Aslam et al., 2019, Luo et al., 2020). Under shade, PIFs recruit the INO80 complex to shade responsive genes and promote H2A.Z eviction (Willige et al., 2021). The interaction is usually mediated by EIN6 ENHANCER (EEN), but EEN-independent mechanisms for H2A.Z removal also exist. INO80 was also found to repress light-induced genes, including *HY5* (Yang et al., 2020). Moreover, PIF7 also recruits histone chaperone ANTI-SILENCING FACTOR 1 (ASF1) and HISTONE REGULATOR HOMOLOG A (HIRA) under shade, and establishes a PIF7-ASF1-HIRA regulatory module, involved in increasing the H3.3 levels on a subset of actively transcribed shade-induced genes (Yang et al., 2023). Collectively this suggests that PIFs and particularly PIF7 control transcriptional output in response to shade not only by direct activation or repression of gene transcription, but also through remodeling of the chromatin landscape of its target genes.

This led us to hypothesize that the local chromatin environment of PIF binding sites might also have a role in controlling shade triggered genome reprogramming. In this study we have combined RNA-seq and Assay for Transposase Accessible Chromatin using sequencing (ATAC-seq) to determine if the PIF-regulated transcriptional response to LRFR depends on chromatin accessibility. Using ATAC-seq, transcription data, and ChIP-seq data, we observe changes in chromatin accessibility of a set of PIF-regulated genes. Our data indicate that the main driving force by which PIFs regulate gene expression is through increased occupancy of their binding sites, without remodeling of the chromatin accessibility for most of their binding sites.

## RESULTS

### PIF7 binding sites are accessible in HRFR

To investigate the connection between the transcriptional response to neighbor proximity (using a LRFR light treatment) and chromatin accessibility, we performed ATAC-seq and RNA-seq. We exposed the seedlings to short (1h) and long (25h) LRFR treatments (Figure 1A) in which neighbor proximity was simulated by supplementing white light with far-red light. We detected 21303 accessible chromatin sites, ∼90% of which are present in regions less than 2 kb 5’ of the TSS, hereafter called promoters (Supplementary Figure 1A). These accessible chromatin regions mostly lack DNA methylation, as described for seedlings grown under similar 16 h-light/8 h-dark cycles (Supplementary Figure 1B) (Zhou et al., 2022), suggesting that prior to a LRFR stimulus (in High R/FR abbreviated HRFR), chromatin in gene regulatory regions is generally accessible for the transcriptional machinery and transcription factors.

**Figure 1.**
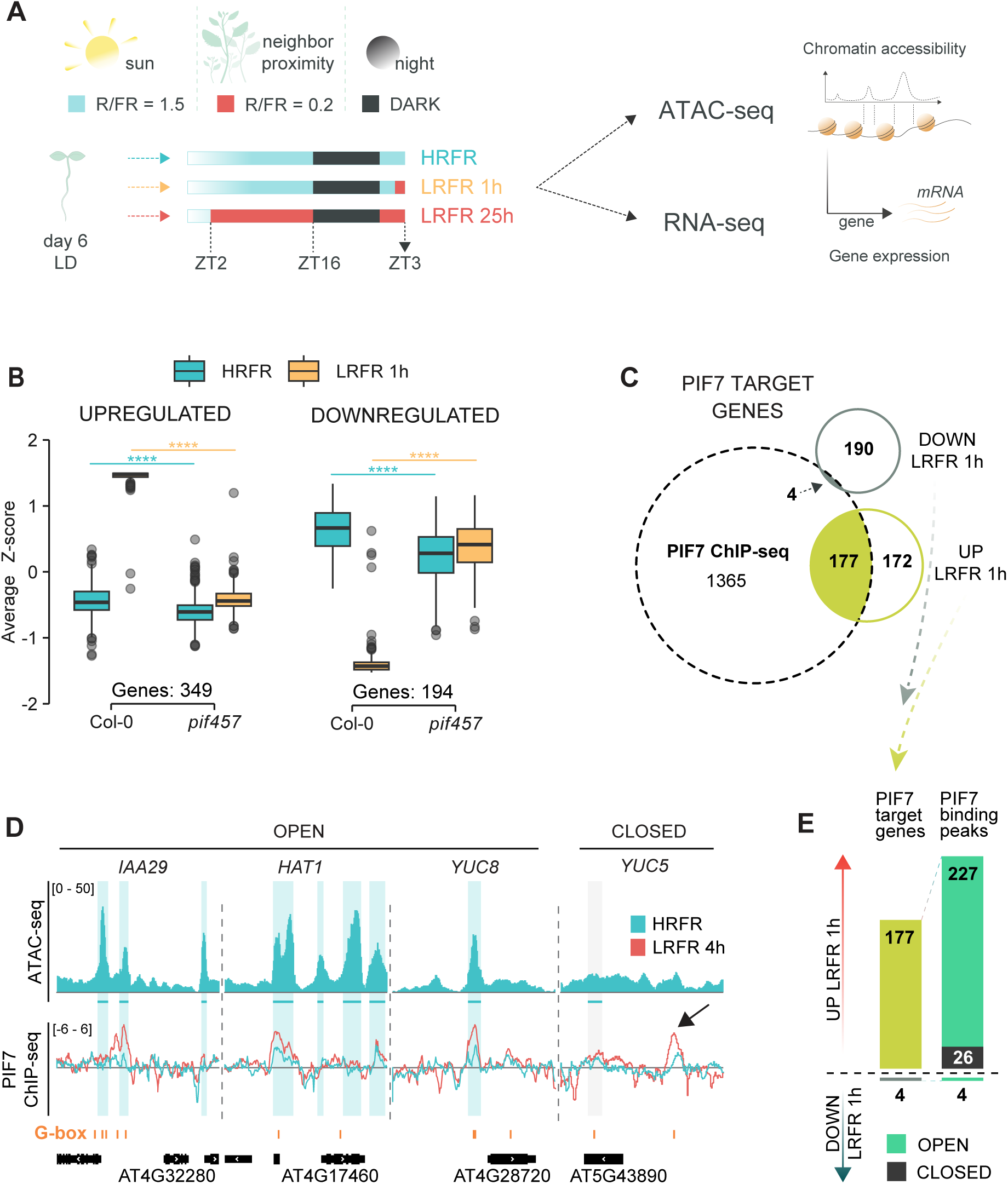
PIF7 binding sites are accessible in HRFR. A. Experimental set-up for INTACT/ATAC-seq and RNA-seq. Seedlings were grown either in HRFR for 7 days (HRFR), moved to LRFR for 1h at ZT2 of day 7 (LRFR 1h) or moved to LRFR at ZT2 of day 6 until day 7 (LRFR 25h). Samples were collected at ZT3 on day 7 and processed for ATAC-seq or RNA-seq. B. PIF457 dependent and shade regulated genes upon exposure to 1h of LRFR from RNA-seq in comparison of Col-0 to *pif457* (padj < 0.05, log2FC – no cutoff). C. PIF7 target genes defined from the overlap of PIF7 binding sites (4h of LRFR exposure) (Willige et al., 2021) and PIF457 dependent DEGs at LRFR 1h (padj < 0.05). Seedlings were grown as in A. D. IGV view of selected PIF7 target genes in HRFR. Upper panel shows ATAC-seq, lower panel shows PIF7 ChIP-seq. Three tracks of biological replicates of ATAC-seq are averaged. One track of PIF7 ChIP-seq for HRFR and 4h of LRFR exposure is shown (Willige et al., 2021). E. Number of binding peaks of PIF7 target genes located on accessible chromatin regions.

Since PIF7 is a major transcription factor regulating LRFR responses, with contributions from PIF4 and PIF5 (Li et al., 2012, de Wit et al., 2016b) we wanted to specifically assess the chromatin environment of PIF7 binding sites. Therefore, we restricted the analysis to a set of shade-regulated genes bound by PIF7 in LRFR. For this purpose, we reanalyzed a published PIF7 ChIP-seq dataset (Willige et al., 2021) and compared it to PIF457-regulated genes in response to short term LRFR (Figure 1B). In response to 1h of LRFR, 349 genes were upregulated, and 194 genes were downregulated in Col-0 compared to *pif457* (Figure 1B, Supplemental table 2). These acute responding genes comprise most known shade marker genes, including *PIL1*, *ATHB2*, and *HFR1*. We, therefore, defined potential PIF7 targets as genes bound by PIF7 within 3 kb upstream of the TSS or 1 kb downstream of the TES in response to 4h of LRFR (Willige et al., 2021) and identified 1546 genes (Figure 1C). By intersecting these lists, we defined 177 shade-upregulated and 4 -downregulated genes as direct PIF7 targets (Figure 1C, Supplemental table 3). The majority of PIF7 binding peaks are found on accessible chromatin regions (open) in seedlings grown in HRFR, with only around 10% of them located in non-accessible regions (closed) (Figure 1D and E), suggesting that PIF7 can easily access its target binding sites prior to a LRFR cue.

### PIFs promote transcriptional response to shade through their increased accumulation and gene occupancy

Strong and fast initial regulation of shade responsive genes is followed by an attenuation of this response under prolonged shade (Sessa et al., 2005, de Wit et al., 2015, Kohnen et al., 2016). The molecular/regulatory mechanisms underlying this PIF-dependent temporal regulation of transcription are, however, not entirely understood. We, therefore, investigated what may explain this transcription pattern induced by PIFs. Our RNA-seq analysis identified 1148 unique genes that were differentially expressed in Col-0 in response to LRFR exposure for 1 h and/or 25 h (Figure 2A, Supplementary Figure 2A and B, Supplemental table 4). Clustering analysis of these differentially expressed genes identified 2 clusters (1 and 2) in which the transient upregulation observed in Col-0 was largely abolished in the *pif457* triple mutant (Figure 2A). These 2 clusters are enriched in terms such as shade avoidance and auxin response (Supplemental table 4), in accordance with the prominent role of these three PIFs in shade responses (de Wit et al., 2016b, Ince et al., 2022). A similar transient pattern is observed with downregulated genes of cluster 7 (Figure 2A). In addition, we observed many genes that respond more slowly to LRFR (clusters 3, 4, and 5) and whose expression, with the notable exception of cluster 4, was largely dependent on PIF457. Also, we find a significant enrichment related to flavonoid biosynthesis and metabolism in clusters 3 and 4 (Supplemental table 4).

**Figure 2.**
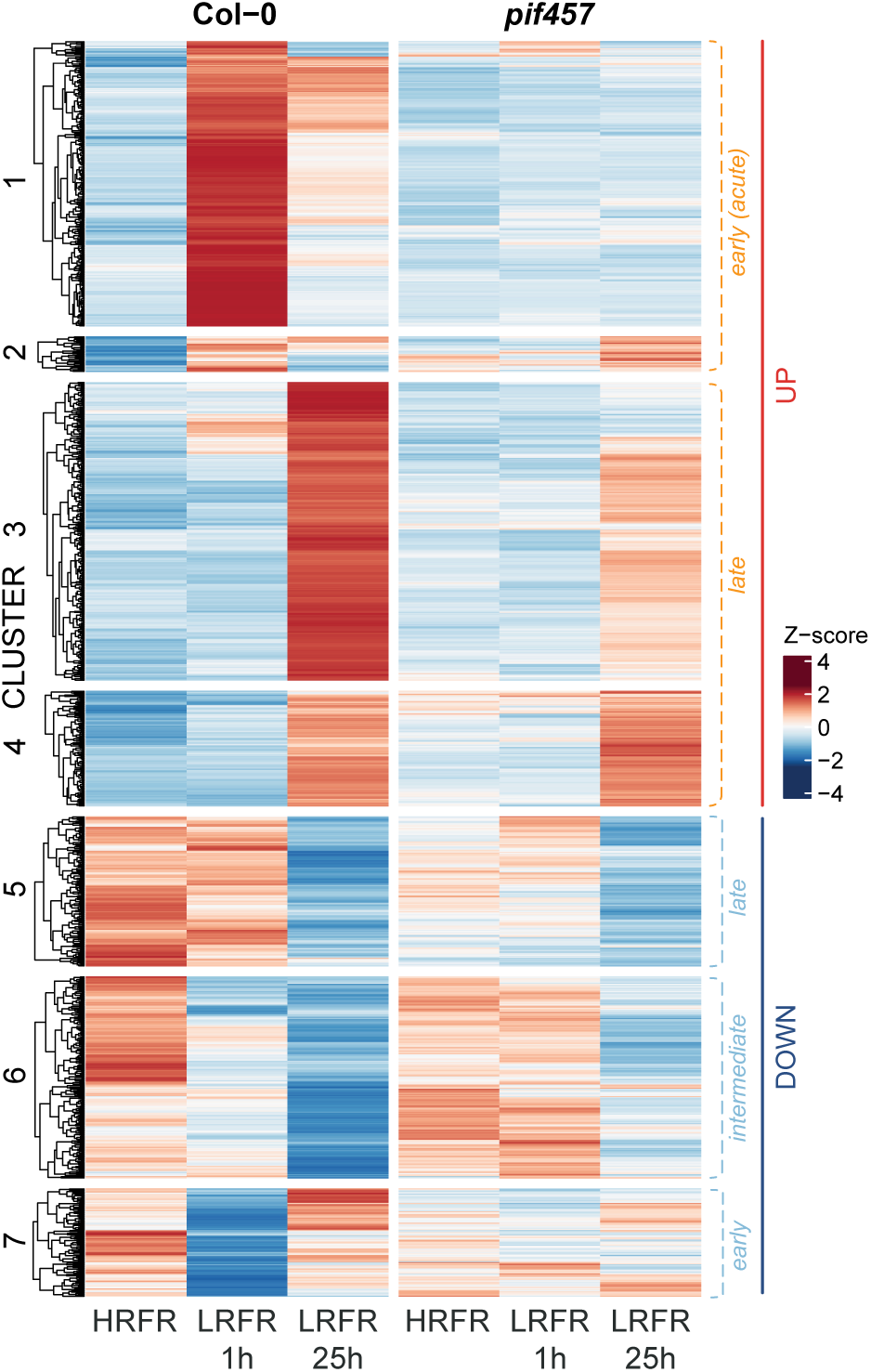
PIFs promote transcriptional response to shade. Heatmap of differentially expressed genes (DEGs) in HRFR, 1h and 25h of LRFR in Col-0 and *pif457* mutant. (padj < 0.05, abs(log2FC) > 0.6).

Since active phyB regulates the turnover of PIFs and reduces their activity (Leivar et al., 2020, Ni et al., 2013, Shen et al., 2008, Shen et al., 2007, Park et al., 2018, Park et al., 2012, Yoo et al., 2021, Xie et al., 2023, Li et al., 2012, Willige et al., 2021), we checked the expression and protein levels of PIF4, PIF5, and PIF7 to determine whether PIF levels may underlie the pattern of PIF target gene expression. These PIFs are known to be under strong circadian clock regulation, with a peak in expression around ZT3 and highest protein accumulation at midday in LD conditions (Galvao et al., 2019, Nozue et al., 2007). In our conditions, the expression of *PIF4* and *PIF5* was shade-unresponsive, while *PIF7* expression was reduced in response to 1 h and 25 h of LRFR (Figure 3A), possibly as a compensatory mechanism of the shade avoidance response. Unlike gene expression, PIF4 and PIF5 protein levels significantly increased transiently in response to shade (Figure 3A), followed by a return to HRFR levels after prolonged LRFR exposure (Figure 3A). PIF7 total protein levels were overall more stable, and only a mild and non-significant increase of PIF7 levels was seen at 1 h of LRFR (Figure 3A). A somewhat distinct LRFR regulation of PIF7 from the one of PIF4 and PIF5 has been suggested, involving regulation of its phosphorylation status and phyB sequestration into condensates (Li et al., 2012, Leivar et al., 2020, Xie et al., 2023). Next, we examined recruitment of PIFs to their binding sites using ChIP-qPCR. We tested PIF binding to canonical CACGTG motifs (G-box) upstream of the *PIL1* TSS to determine whether there was a correlation between PIF protein levels and recruitment to chromatin in response to shade. We observed a strong correlation between PIF4 protein levels (Figure 3A) and *PIL1* promoter occupancy (Figure 3B), and a similar behavior was seen with PIF7 (Figure 3B). The same trend was observed for other PIF-regulated genes (Supplementary Figure 3 A-B), indicating that a transient increase in PIF protein levels and/or overall stability of PIFs led to higher occupancy of their binding sites, providing a possible explanation for the transient upregulation of many early PIF target genes (Figure 3C).

**Figure 3.**
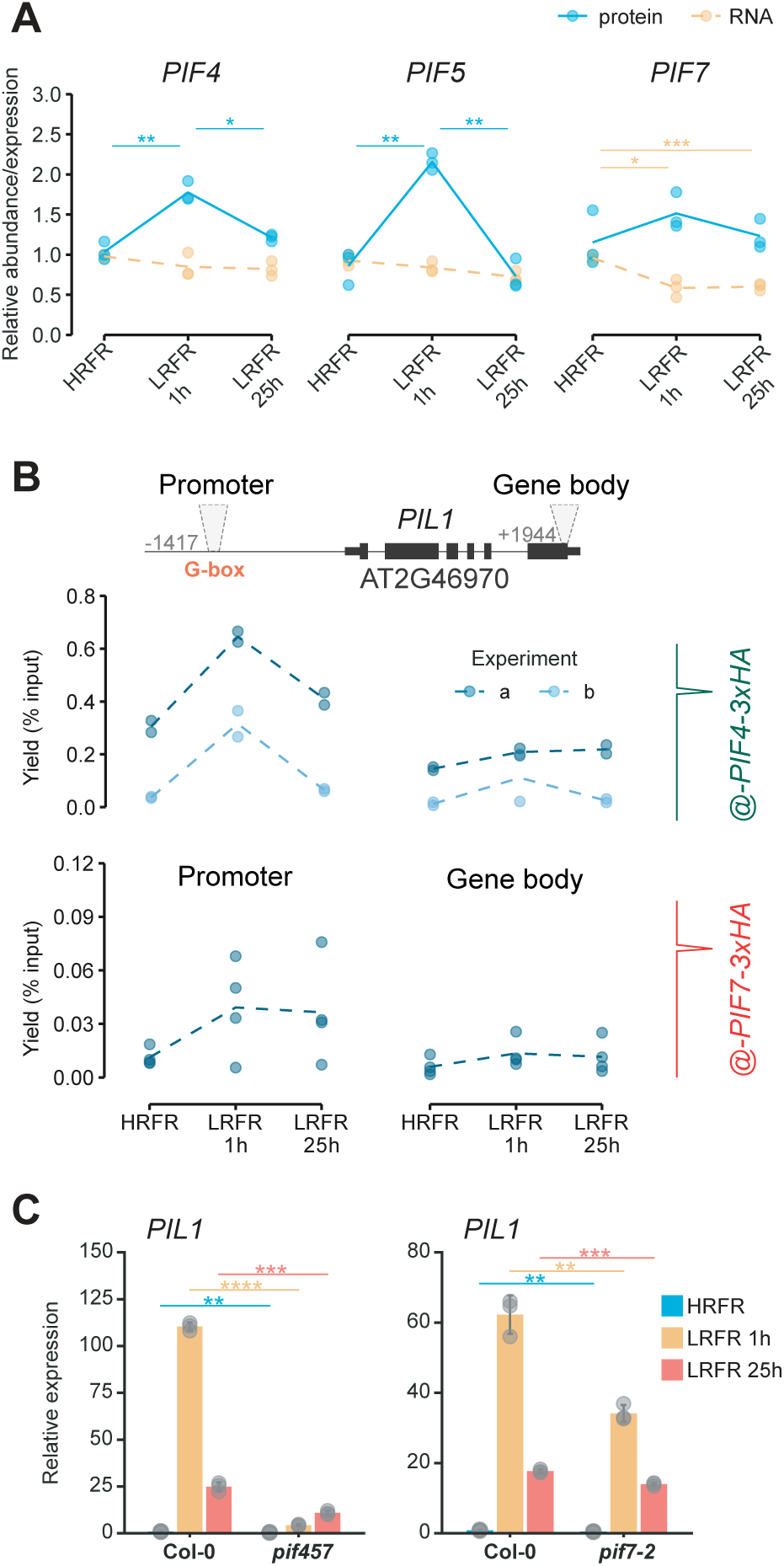
Transient increase in accumulation of PIFs correlates with gene occupancy in response to LRFR. A. Relative gene expression (yellow) of *PIF4*, *PIF5* and *PIF7* normalized to *YLS8* and *UBC* levels. Relative protein abundance (blue) of PIF4, PIF5 and PIF7. PIF4 protein levels were normalized to tubulin levels, while PIF5 and PIF7 protein levels were normalized to DET3 levels. Three biological replicates (represented as dots) were used for calculating average values (represented as lines). B. ChIP-qPCR of pPIF4:PIF4-3xHA line (in pif4-101) for *PIL1* locus (upper panel, two biological replicates from two independent experiments are presented). Lower panel displays ChIP-qPCR of four biological replicates of pPIF7:PIF7-3xHA line (in *pif7-2*) for *PIL1* locus. C. Relative expression of *PIL1* in *pif457* (left panel) and *pif7-2* (right panel) mutants. Seedlings were grown either in HRFR for 7 days (HRFR), moved to LRFR for 1h at ZT2 of day 7 (LRFR 1h) or moved to LRFR for at ZT2 of day 6 until day 7 (LRFR 25h). Samples were collected at ZT3 on day 7. Asterisks represent statistical significance (Students T-test, *p<0.05, **p<0.01, ***p<0.001, ****p<0.0001).

### Low R/RF promotes dynamic changes of accessible chromatin regions

To examine the effect of LRFR on accessible chromatin regions we performed differential accessibility analysis of ATAC-seq. In total, we detected 84 differentially accessible regions (DARs), 32 in response to 1 h of LRFR (Figure 4A, Supplementary Figure 4A, Supplemental table 5), and 61 in response to 25 h of LRFR (Figure 4A, Supplementary Figure 4A, Supplemental table 5). Under short LRFR exposure, the regions were prevalently becoming more accessible compared to HRFR, with less accessible regions appearing after long LRFR exposure (Figure 4A, Supplementary Figure 4A). Genes associated with DARs that were not expressed in Col-0 were excluded from further analysis, leaving 72 DARs associated with 65 genes (Supplemental table 5). In general, these DARs were mostly found within 0.1-2 kb 5’ of the TSS (33) and within +/- 0.1 kb of the TSS (25), with 17 DARs directly bound by PIF7 (Figure 4B, Supplemental table 5).

**Figure 4.**
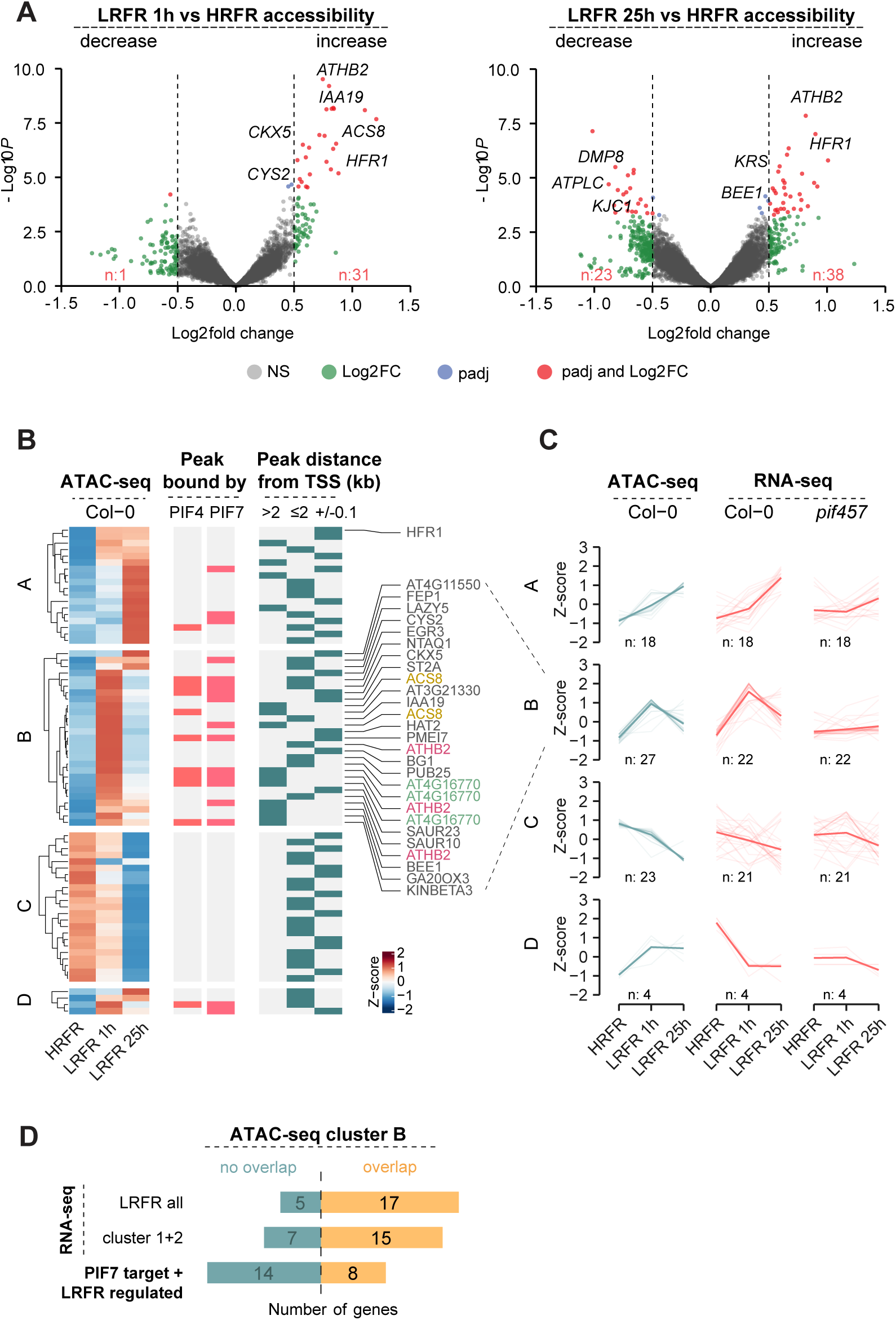
Chromatin accessibility in response to LRFR affects a moderate number of genes. A. Volcano plots of differentially accessible regions (DARs) in comparisons of 1h of LRFR (left panel) and 25h of LRFR (right panel) versus HRFR in Col-0. Statistical cutoff is set at padj < 0.05 and abs(log2FC) > 0.5. B. Heatmap of differentially accessible regions (DARs) in Col-0 in HRFR, LRFR 1h and LRFR 25h, hierarchically clustered into four distinct clusters based on ATAC-seq and RNA-seq (left panel). DARs bound by PIF4 (Pfeiffer et al., 2014) and PIF7 (Willige et al., 2021) are represented in the middle panel. The distance of the DAR from TSS is indicated on right panel. C. ATAC-seq counts of DARs in Col-0 and the expression of genes in Col-0 and *pif457* associated with the DARs are represented as an average z-score. Thick line is the average trend line. The number of DARs and genes in each cluster is displayed below the line plots. D. Number of genes from ATAC-seq cluster B that overlap with all DEGs from RNA-seq or only with acute shade responding clusters 1 and 2 and with PIF7 and shade regulated target genes

We performed hierarchical clustering of the 72 DARs and divided them into four distinct clusters, considering chromatin accessibility and gene expression of the associated genes in Col-0 (Figure 4B). Cluster A consists of DARs that increased accessibility after 25 hours of LRFR exposure, with only a few of these sites bound by PIF4 or PIF7 (Figure 4B), as defined based on the data from (Willige et al., 2021, Pfeiffer et al., 2014). In contrast, cluster B contains DARs that transiently increased accessibility after 1 h of LRFR, and many of these are bound by PIF4 and PIF7 (Figure 4B). Importantly, the chromatin accessibility pattern of cluster B reflected the transcriptional response of Col-0 and was dependent on PIF457 (Figure 4C). Most genes in cluster B were regulated by LRFR (Figure 4D) and included direct PIF7 targets, such as *ATHB2*, *ACS8*, *IAA19*, and *HAT2* (Supplementary Figure 4C). Gene Ontology (GO) enrichment analysis of this cluster confirmed a significant enrichment in terms related to the auxin response and shade avoidance (Supplementary Figure 4D). While transcriptional and accessibility patterns closely matched in clusters A and B, the same cannot be said for cluster C, which were more variable or cluster D, where accessibility increased yet transcription decreased in response to LRFR (Figure 4C). We note that while for some shade-induced genes, we saw a chromatin accessibility pattern that correlated with gene expression, this group represents less than 5% of transiently shade-induced genes from RNA-seq clusters 1 and 2 (Supplemental table 4).

Since auxin was found to control chromatin accessibility during developmental reprogramming (Wu et al., 2015, Wang et al., 2020, Wu et al., 2022), we hypothesized that chromatin accessibility might also be affected by the shade-induced increase in auxin levels. Because SAV3 mediates a critical auxin biosynthetic step in the formation of indole-3-pyruvic acid (IPA) (Tao et al., 2008), we examined the *sav3-2* mutant by ATAC-seq and compared it to Col-0. The most prominent shade responsive genes such as *ATHB2* and *HFR1*, which are SAV3 independent (Li et al., 2012) did not significantly differ between *sav3-2* and Col-0 (Supplemental table 5). However, we observed a weaker response to short term shade for genes present in the PIF dependent cluster B (Supplementary Figure 5A-B). PIF independent clusters showed similar patterns of chromatin accessibility changes in Col-0 and *sav3-2* (Supplementary Figure 5A-B). Yet, the reduced response of cluster B in *sav3-2* suggests a partial dependence on auxin. However, as observed in Col-0, this pattern of chromatin accessibility was only seen for a subset of shade-regulated genes.

In addition to transcriptional changes LRFR promotes various epigenetic changes. Some are well described and induced by PIFs, such as the two marks of active transcription H3K9ac and H3K9me3 (Willige et al., 2021, Calderon et al., 2022). We used published ChIP-seq datasets of these two histone modifications (GSE139296, PRJNA839161) to examine our ATAC-seq gene clusters. We found that, in contrast to other clusters, genes in PIF-dependent cluster B are also more enriched in H3K9ac and H3K9me3 under LRFR (Supplementary Figure 6 A-B).

### phyB and PIFs contribute to changes in chromatin accessibility

To investigate whether the phyB-PIF module is involved in regulating chromatin accessibility in response to LRFR we analyzed DARs of two typical shade-upregulated genes *ATHB2* (from cluster B) and *HFR1* (from cluster A) (Figure 5A, Supplementary Figure 7C-D). We chose these genes because they have different expression patterns (*ATHB2* expression is more transiently induced by LRFR, while *HFR1* expression remains high in LRFR) and because their DARs have different characteristics: *ATHB2*’s DARs are 5’ upstream of its TSS and are PIF7 binding sites, while *HFR1*’s DAR is at the TSS and is not a PIF7 binding site. In all three cases, LRFR led to their enhanced accessibility (Figure 5A). Of particular note is that the DAR at *HFR1*’s TSS does not comprise a G-box, while both PIF7 binding sites upstream of *ATHB2* (P1, P2) comprise G-boxes. We compared chromatin accessibility at these DARs with regions on the gene body (G) of both genes, which were less accessible and did not change in response to LRFR (Figure 5A). Using CoP-qPCR, a method to isolate accessible chromatin regions and compare them to the input to test for enrichment (Zhang et al., 2020a), we were able to confirm a significant increase in accessibility of *ATHB2* and *HFR1* DARs in Col-0, validating the ATAC-seq results (Figure 5B). Since these changes were LRFR-mediated, we first decided to test the role of phyB using both over-expressors and a loss-of-function mutant. Overexpression of phyB resulted in a strong repression of hypocotyl elongation in both HRFR and LRFR, while the absence of phyB promoted hypocotyl elongation (Supplementary Figure 7 A-B). In addition, expression of the shade-marker genes *PIL1* and *HFR1* was de-repressed in *phyB-9*, while strongly reduced in the over-expression line (Supplementary Figure 7 C-D). In *phyB-9* mutant, *ATHB2* DARs were already more accessible in HRFR and remained more accessible in all conditions (Figure 5B). The same tendency was observed at the *HFR1* TSS in the *phyB-9* mutant (Figure 5B), although this region is not directly bound by PIFs. In contrast, phyB overexpression prevented both *ATHB2* DARs from increasing their accessibility in response to shade (Figure 5B).

**Figure 5.**
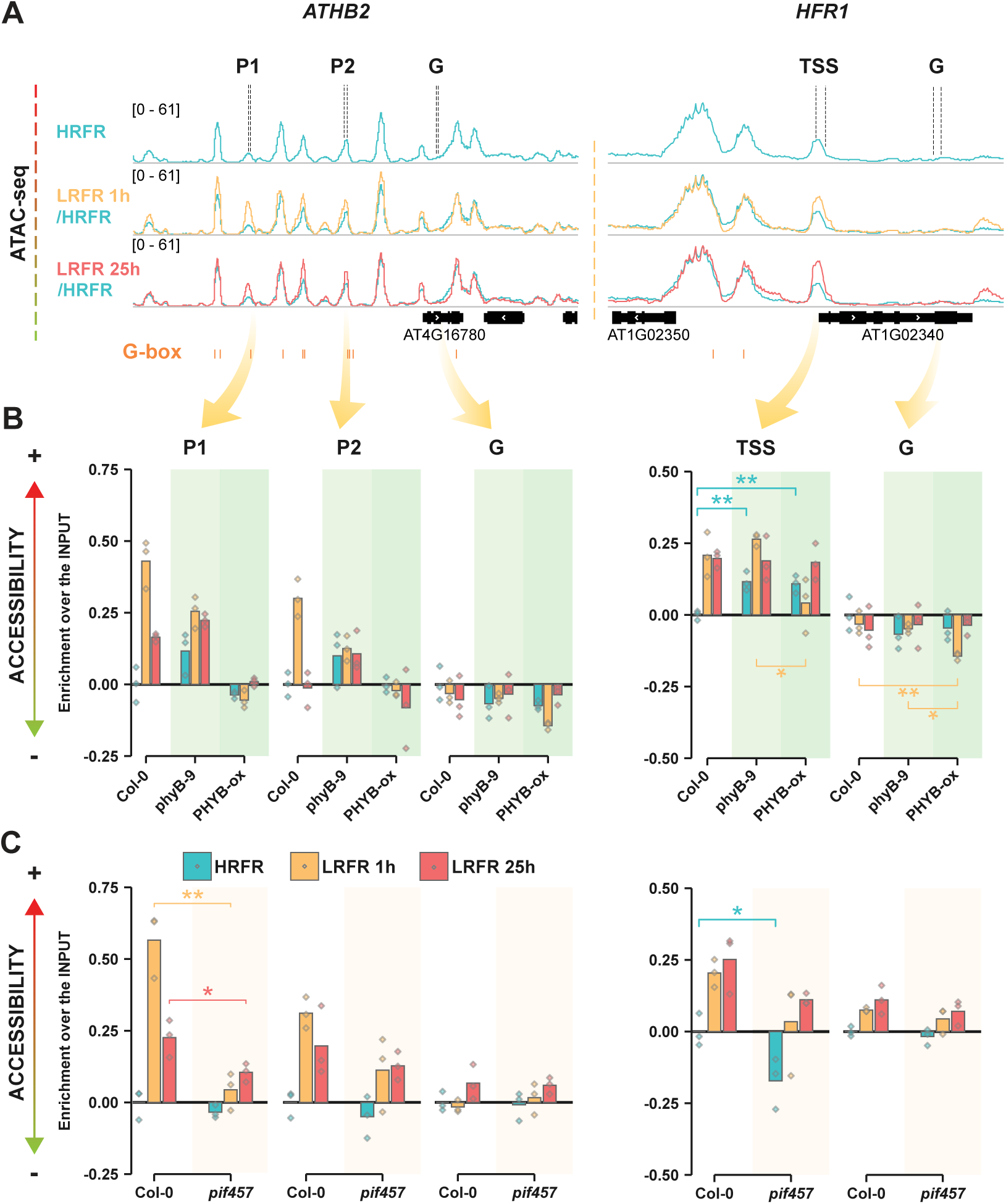
Increase in chromatin accessibility of a set of shade regulated genes is induced by PIFs in response to LRFR. A. IGV view of shade regulated genes *ATHB2* and *HFR1* with changes in chromatin accessibility in response to LRFR in Col-0. Regions tested by CoP-qPCR are marked above the panel. G-boxes are indicated in orange below the panels. ATAC-seq tracks are an average of 3 biological replicates. B. Chromatin accessibility of *ATHB2* and *HFR1* assayed by CoP-qPCR in Col-0, *phyB-S* and PHYB-ox (35S:PHYB-GFP). The accessibility is relative to Col-0 in HRFR (represented as 0). Asterisks represent statistical significance (Tukey HSD, *p<0.05, **p<0.01, ***p<0.001, ****p<0.0001). C. Chromatin accessibility of *ATHB2* and *HFR1* assayed by CoP-qPCR in Col-0 and *pif457*. The accessibility is relative to Col-0 in HRFR (represented as 0). Asterisks represent statistical significance (Tukey HSD, *p<0.05, **p<0.001, ***p<0.001, ****p<0.0001).

Considering PIFs as major targets of phyB regulation and recent findings indicating a role of PIFs in altering the chromatin landscape (Leivar et al., 2020, Ni et al., 2013, Shen et al., 2008, Shen et al., 2007, Xie et al., 2023, Willige et al., 2021, Xue et al., 2021, Zhao et al., 2023, Verma et al., 2024), we hypothesized that the effect of phyB on chromatin accessibility may be achieved through PIFs. We, therefore, determined the role of PIF457 in LRFR-induced hypocotyl elongation, expression of shade marker genes, and DARs by using the *pif457* triple mutant (Supplementary Figure 7). The response of both *ATHB2* DARs to LRFR was strongly reduced in *pif457*, indicating that PIFs are necessary for the increase in their accessibility (Figure 5C). The same effect was seen for *IAA19* (Supplementary Figure 7 E-F). Interestingly, we also observe overall reduced accessibility of *HFR1* TSS in *pif457*, suggesting a strong effect of PIF457 on this region, although the response to LRFR maintained a similar magnitude (Figure 5C).

*HFR1* has a particular transcription pattern that remains at high levels even after 25h of LRFR (Supplementary Figure 8 C-D) (Sessa et al., 2005). This pattern correlates with the increased accessibility of its TSS. To investigate the regulation of *HFR1* TSS accessibility and a potential connection between chromatin accessibility and transcription, we investigated HY5 as a potential candidate. HY5 binds to the TSS of *HFR1* (Burko et al., 2020) and contributes to promoting its expression in response to LRFR (Supplementary Figure 8 C), HY5 also repressed hypocotyl elongation in HRFR and LRFR (Supplementary Figure 8 A). In addition, we investigated the role of HY5 HOMOLOGUE (HYH), which regulates transcriptional reprogramming together with HY5 (Sharma et al., 2023). However, in our conditions, *hy5hyh* did not significantly differ from *hy5* (Supplementary Figure 8 B, D). Finally, we did not observe significant changes in *HFR1* TSS opening in response to long-term LRFR between Col-0 and *hy5*or *hy5hyh* (Supplementary Figure 8 E-F), suggesting that HY5 and HYH do not regulate *HFR1* TSS chromatin accessibility.

## DISCUSSION

Our aim was to determine the potential role of chromatin accessibility during LRFR-induced transcriptional reprogramming. We focused on the roles of PIFs and particularly PIF7, which is known to play a central role in LRFR-induced changes of gene expression (Li et al., 2012, de Wit et al., 2016a, Willige et al., 2021, Cai and Huq, 2024, Favero, 2020). Our data indicates that chromatin accessibility is not a major barrier for PIF7 binding and activity prior to shade treatment, given that PIF7 binding sites were accessible prior to LRFR treatment for approximatively 90% of the putative PIF7 target genes (Figure 1 D-E, Supplementary Figure 1 A-B). With relatively easy access to its binding sites, PIF-induced transcription can be very rapid (Figure 2), consistent with previous publications (Kohnen et al., 2016, Ciolfi et al., 2013, Sessa et al., 2005, Salter et al., 2003, Willige et al., 2021). Transient shade-regulated changes in transcript abundance observed for hundreds of genes are also unlikely to depend on chromatin accessibility. Indeed, for most PIF7-target genes we did not observe changes in chromatin accessibility in response to LRFR (Figure 4, Supplemental table 2-3). Similarly, chromatin accessibility does not seem to explain the distinct response between rapidly LRFR-induced (clusters 1, 2, in Figure 2) and slowly induced genes (cluster 3, in Figure 2). Chromatin accessibility was also analyzed during de-etiolation (Sullivan et al., 2014), a developmental process characterized by altered expression of thousands of genes (Dong et al., 2014). Accessibility changes observed on regulatory regions during de-etiolation are found only for a small subset of differentially expressed genes (Sullivan et al., 2014, Dong et al., 2014), suggesting that genome-wide transcriptional regulation mediated by light may not need extensive remodeling of chromatin accessibility. Nevertheless, we identified a small subset of shade-regulated PIF7 target genes with changes in chromatin accessibility either at PIF binding sites or at the TSS (Figure 4). These include well known shade marker genes such as *ATHB2*, *HFR1* and *IAA19*. Most of these genes are found within a slightly larger cluster B of rapidly upregulated genes in response to LRFR with chromatin accessibility changes along their regulatory regions (Figure 4, Supplementary Figure 1, Supplemental table 5). We found that these genes are not expressed in a specific cell type under HRFR (i.e., the mesophyll as the most abundant cell type) (Han et al., 2023), but they are characterized by two marks of active transcription, H3K9ac and H3K4me3 (Supplementary Figure 6), that are prevalent in PIF regulated genes in response to LRFR (Calderon et al., 2022, Willige et al., 2021).

In our study LRFR triggers moderate but significant changes in chromatin accessibility (Figure 5). To establish what regulates these changes, we investigated auxin as the major driving force of physiological adaptations to shade. Though auxin-related genes show strong transcriptional response under shade (Tao et al., 2008, Kohnen et al., 2016), the reduction in chromatin accessibility of a set of genes in PIF regulated cluster B observed in *sav3-2* is moderate (Supplementary Figure 5). One hypothesis is that the transient auxin increase during early perception of neighbor threat (Pucciariello et al., 2018) is not capable of inducing profound remodeling of chromatin accessibility as seen during developmental reprogramming (Wu et al., 2015, Wang et al., 2020, Wu et al., 2022). However, given that four genes from cluster B with reduced changes in chromatin accessibility in *sav3-2* also show reduced LRFR-induced expression in this mutant (Li et al., 2012), this suggests a link between changes in chromatin accessibility and increased expression in this subset of genes.

Chromatin accessibility at most PIF7 binding sites is stable (Figure 1), but a group of LRFR regulated genes displays rather dynamic changes at their PIF4/PIF7 binding sites (Figure 4). Consistent with the role of phyB as the critical regulator of PIFs activity (Leivar et al., 2020, Ni et al., 2013, Shen et al., 2008, Shen et al., 2007, Park et al., 2018, Park et al., 2012, Yoo et al., 2021, Xie et al., 2023), we found that PIF7 bound DARs showed increased chromatin accessibility in *phyB-9*, in contrast to overexpressing phyB, supporting the role for PIFs in regulating chromatin accessibility in response to shade (Figure 5). Whether PIFs can directly remodel the chromatin landscape of the genes they regulate was not fully investigated. However, several publications indicate that PIF-mediated transcription is accompanied by chromatin changes of the transcribed regions (Willige et al., 2021, Xue et al., 2021, Zhao et al., 2023, Verma et al., 2024). In favor of PIFs ability to promote nucleosome remodeling are the described lower chromatin accessibility found at the promoters of *BBX21* and *GLK1* genes in the *pifQ* mutant under darkness (Guo et al., 2023) and the capacity of the PIF4 and PIF7 to directly interact with the INO80 complex to promote H2A.Z eviction (Xue et al., 2021, Willige et al., 2021, Verma et al., 2024, Zhao et al., 2023). Moreover, PIF7 recruits the INO80 complex in response to shade at transcribed genes (Willige et al., 2021). We can speculate that the remodeling of the chromatin accessibility found on PIF binding sites triggered by PIFs might be a consequence of a similar recruited complex of the SWI/SNF type. Importantly, we observe a similar trend in chromatin accessibility and transcriptional changes, and for cluster B this seems to correlate with the occupancy of PIF binding sites. Although we are not able to concurrently assess whether differences in the chromatin landscape of PIF binding sites might be associated with the observed transcriptional pattern differences, Willige et al. notice increased hyperacetylation around DNA regulatory regions on the promoters of some shade responsive genes such as *ATHB2* and *HAT3* (Willige et al., 2021). This hyperacetylation seems to appear faster than on the gene bodies, even though on much smaller scale (Willige et al., 2021). To what extent these changes influence transcriptional response to LRFR needs more attention, as some chromatin modifications occur rapidly (Willige et al., 2021), while others are trailing marks of active PIF-mediated transcription (Calderon et al., 2022).

Based on our study, we suggest that the chromatin accessibility of PIF binding sites is not a limiting factor in the regulation of gene transcription in response to LRFR, as the sites are easily accessible prior to LRFR exposure. Increase in PIF protein levels and gene occupancy seem to be the key drivers of this transcriptional response (Figure 2 and 3 A-C). Several other factors probably contribute to the dynamic regulation of shade responses, including regulation of phosphorylation status of PIF7 (Li et al., 2012, Leivar et al., 2020). Moreover, PIFs are tightly regulated by the circadian clock (Soy et al., 2016, Zhang et al., 2020b) and modulated by interaction with factors such as HFR1, DELLAs and HY5 (de Wit et al., 2016b, Xu et al., 2017, Cai and Huq, 2024, Favero, 2020, Yang et al., 2021). Besides, while HY5 contributes to the transcriptional regulation of *HFR1*, it does not appear to affect HFR1 TSS chromatin accessibility (Supplementary Figure 8), suggesting a primary role for PIFs. More genome wide studies focused on specific cell types are needed to assess PIF7 mediated chromatin remodeling, particularly around critical DNA regulatory motifs.

## CONCLUSIONS

Our study highlights how PIFs primarily regulate transcriptional responses to shade by increasing their occupancy at already accessible chromatin regions, with a moderate remodeling of that chromatin landscape. We propose that shade-mediated transcriptional regulation may not require extensive remodeling of DNA accessibility and is instead confined to a small subset of genes. These genes might share specific genomic features that we are currently unable to identify. While histone modifications of PIF regulated genes are significant indicator of their transcriptional activity, the main regulatory force of PIF-induced gene regulation appears to be PIF protein dynamics and their interaction with the pre-existing accessible chromatin landscape.

## MATERIALS AND METHODS

### Plant material

*Arabidopsis thaliana* pPIF7:PIF7-3xHA (in *pif7-2*), pPIF4:PIF4-3xHA (in *pif4-101*) and p35S:PHYB-GFP (PHYBox, in *phyB-9*) lines and *pif7-2*, *pif4-101 pif5-3 pif7-1* (*pif457*), *phyB*-*9*, *sav3-2*, *hy5* and *hy5hyh* mutants been previously described (Galvao et al., 2019, Zhang et al., 2017, Leivar et al., 2008, de Wit et al., 2015, Neff and Chory, 1998, Oyama et al., 1997, Tao et al., 2008, Zoulias et al., 2020, Medzihradszky et al., 2013).

### Growth conditions and light treatments

The Arabidopsis thaliana seedlings used in this study, except for hypocotyl elongation, were grown on ½ MS plates (without sucrose) for 7 days at 21°C under 16h light/8h dark conditions (high Red/Far-Red light ratio - HRFR) or treated with low Red/Far-Red light (LRFR). Seedlings were treated with LRFR from ZT2 of day 6 until ZT3 of day 7 (LRFR 25h) or from ZT2 until ZT3 of day 7 (LRFR 1h). HRFR ratio corresponds to approximately 1.5 and a total fluence rate of PAR ∼47-50 µmol/m^-2^s^-1^. The ratio of LRFR treatment was from 0.13-0.2. For hypocotyl elongation experiments, seedlings were grown on ½ MS plates (without sucrose) for 7 days at 21°C in HRFR or treated with LRFR from day 4 until day 7. Plants for seed production were grown on soil under 16h light/8h dark conditions in walk-in growth chambers.

### Generation of transgenic lines for INTACT-ATAC

For the INTACT-ATAC, the lines expressing both the pUBQ10::BirA and pUBQ10::NTF transgenes were generated in WT Col-0 and introduced into the *sav3-2* mutant by crossing. Briefly, pUBQ10::BirA was generated as described in (You et al., 2017). The pUBQ10:NTF construct was generated, as also described in (You et al., 2017), cloning the WPP domain of the Arabidopsis thaliana RAN GTPASE ACTIVATING PROTEIN 1 (RanGAP1; At3g63130) at the N terminus, followed by the enhanced GFP protein (eGFP) and the biotin ligase recognition peptide downstream of the UBQ10 promoter. After crossing with *sav3-2* mutant, plants bearing both the pUBQ10:BirA and pUBQ10:NTF were selected for the resistance to glufosinate (BASTA) and kanamycin, respectively. Moreover, the expression of the biotinylated NTF proteins was tested running total protein extracted from homozygous lines on 4-15% gel, transferred on nitrocellulose membrane and probed with anti-Streptavidin antibody conjugated with HRP for chemiluminescent visualization.

### Hypocotyl measurements

Hypocotyl elongation measurements were done as described previously (de Wit et al., 2015). In brief, seedlings were grown on ½ MS vertical plates in a growth incubator in HRFR until day 4 (ZT2) and images were taken. Plates with seedlings were transferred to LRFR or kept in HRFR until taking images on day 7 (ZT2). Hypocotyl length was measured with a MATLAB script developed in the C.F. laboratory.

### Reverse Transcription - quantitative PCR (RT-qPCR)

Total Arabidopsis RNA was extracted using Plant RNeasy kit (Qiagen, Cat. No./ID: 74904) according to the manufacturer’s instructions. cDNA was synthesized using Superscript II Reverse Transcriptase (Invitrogen, Life Technologies) with random oligonucleotides. Quantitative real-time PCR (qPCR) was done in three biological replicates with three technical replicates on QuantStudio 6 Flex Real-Time PCR System (Applied Biosystems). Gene expression data was normalized against UBC and YSL8 genes. Primers used for the qPCR reactions can be found in Supplementary Table 1.

### ChIP-qPCR

For one biological replicate, 15 mg of seeds were sown on ½ MS. Seedlings were harvested, frozen in liquid nitrogen and ground with mortar and pestle. Ground powder was transferred to 10 mL of cold EB1 (60 mM HEPES pH 8, 0.4 M sucrose, 10 mM KCl, 10 mM MgCl_2_, 5 mM EDTA, cOmplete™ Mini EDTA-free Protease Inhibitor Cocktail) supplemented with 1% formaldehyde and incubated for 10 min at RT on a rotating wheel. Crosslinking was stopped by adding freshly prepared 2 M glycine to a final concentration of 0.125 M, incubated for 10 min at RT on a rotating wheel. Extracts were filtered through a layer of Miracloth, centrifuged for 10 min at 4000xg at 4°C and supernatant was removed. Nuclei pellets were resuspended first in 1 mL of EB2 (0.25 M sucrose, 10 mM Tris-HCl pH 8, 1% Triton, 10 mM MgCl_2_, 5 mM beta-mercaptoethanol, cOmplete™ Mini EDTA-free Protease Inhibitor Cocktail), centrifuged for 10 min at 5000xg at 4°C and then resuspended in 300 µL EB3 (1.7 M sucrose, 10 mM Tris-HCl pH 8, 0.15% Triton, 2 mM MgCl_2_, 5 mM beta-mercaptoethanol, cOmplete™ Mini EDTA-free Protease Inhibitor Cocktail). After centrifugation for 1h at 16000xg at 4°C, nuclei pellets were resuspended in 200 µL of NLB (50 mM Tris-HCl pH 8, 10 mM EDTA, 1% SDS, cOmplete™ Mini EDTA-free Protease Inhibitor Cocktail) and sonicated for 15 min on high power (PIF7) and 20 min on low power (PIF4) (SONICATION Bioruptor Digenode: 30s ON 30s OFF, 10 cycles).

The chromatin was immunoprecipitated with mouse monoclonal HA-Tag Antibody (F-7) (Santa Cruz Biotechnology, Cat No./ID: sc-7392) coupled to Dynabeads™ Protein A and Protein G mixture (Invitrogen, Cat No./ID: 10001D and 10003D). Chromatin was eluted in EB (1% SDS, 0.1 M NaHCO3) at 65°C for 15 min twice. Input and IP were reverse crosslinked overnight at 65°C with NaCl (0.2 M final) then treated with RNase A (Qiagen, Cat No./ID: 19101) for 30 min at 37°C, followed by Proteinase K (Fisher Scientific, Cat No./ID: AM2546) for 30 min at 55°C. DNA was purified with QIAquick PCR Purification Kit (Qiagen, Cat No./ID: 28104).

### Western blot - SDS-Page

For protein extractions, 20-25 seedlings were frozen in liquid nitrogen, ground to powder and resuspended in 2x Laemmli buffer (0.125 M Tris-HCl pH 6.8, 4% SDS, 20% glycerol, 10% beta-mercaptoethanol, bromophenol blue). Extracts were heated for 5 min at 95°C, spined down and loaded to 4–15% Mini-PROTEAN® TGX™ Precast Protein Gels (Bio-Rad, Cat No./ID: 4561086). After semi-dry transfer, the membranes were probed with rat monoclonal anti-HA antibody coupled to HRP (Roche, Cat No./ID: 12013819001). Rabbit polyclonal anti-DET3 (Schumacher et al. 1999) and mouse monoclonal anti-tubulin (Abiocode, Cat No./ID: M0267-1a) antibodies were used for normalization. Chemiluminescence was generated with Immobilon Western Chemiluminescent HRP Substrate on Fujifilm ImageQuant LAS 4000 mini-CCD camera system (GE Healthcare).

### CoP (column purified isolation of regulatory elements)

Accessible chromatin was extracted using a modified CoP method (Zhang et al., 2020a). For one biological replicate, 6-10 mg of seeds were sown on ½ MS. Seedlings were harvested, frozen in liquid nitrogen and ground with mortar and pestle. Ground powder was transferred to 35 mL of cold EB1 (60 mM HEPES pH 8, 1 M sucrose, 5 mM KCl, 5 mM MgCl_2_, 5 mM EDTA, 0.6% Triton-X100, cOmplete™ Mini EDTA-free Protease Inhibitor Cocktail) and incubated with 1% formaldehyde for 10 min at RT on a rotating wheel. Crosslinking was stopped by adding 2.4 mL of freshly prepared 2 M glycine, incubated for 10 min at RT on a rotating wheel. Extracts were filtered through a layer of Miracloth, centrifuged for 20 min at 4000xg at 4°C and supernatant was removed. Nuclei pellets were resuspended first in 1 mL of EB2 (0.25 M sucrose, 10 mM Tris-HCl pH 8, 1% Triton, 10 mM MgCl_2_, 5 mM beta-mercaptoethanol, cOmplete™ Mini EDTA-free Protease Inhibitor Cocktail), centrifuged for 10 min at 12000xg at 4°C and then resuspended in 300 µL EB3 (1.7 M sucrose, 10 mM Tris-HCl pH 8, 0.15% Triton, 2 mM MgCl_2_, 5 mM beta-mercaptoethanol, cOmplete™ Mini EDTA-free Protease Inhibitor Cocktail). After centrifugation for 1h at 16000xg at 4°C, nuclei pellets were resuspended in NLB (50 mM Tris-HCl pH 8, 10 mM EDTA, 1% SDS, cOmplete™ Mini EDTA-free Protease Inhibitor Cocktail) and sonicated for 10 min on high power (SONICATION Bioruptor Digenode: 30s ON 30s OFF, 10 cycles).

10% of chromatin was spared as input, the rest was purified using silica membrane columns from Wizard® SV Gel and PCR Clean-Up kit (Promega, Cat No./ID: A9282). Input was treated with RNase A (Qiagen, Cat No./ID: 19101) for 30 min at 37°C, followed with Proteinase K (Fisher Scientific, Cat No./ID: AM2546) for 30 min at 55°C, then at 65°C overnight. DNA was purified with Wizard® SV Gel and PCR Clean-Up kit.

### INTACT/ATAC-seq

For each biological replicate of ATAC-seq, 10 mg of seeds were sown on ½ MS plates to obtain around 0.4 g of seedlings. ATAC-seq was performed as described previously (Bajic et al., 2018) with minor modifications. In brief, the tissue was frozen in liquid nitrogen for nuclei isolation using the INTACT method (You et al., 2017). The frozen tissue was ground with mortar and pestle then transferred to 15 mL tube with NPB (MOPS pH7 20 mM, NaCl 40 mM, KCl 90 mM, EDTA 2 mM, EGTA 0.5 mM, Spermidine 0.5 mM, Spermine 0.2 mM, 0.5 x Complete Protease Inhibitors) and filtered through one layer of Miracloth and 30 µm filter-tubes (Sysmex, Cat No./ID: 04-004-2326). After centrifugation at 1000xg for 10 min at 4°C, the nuclei were resuspended in 1 mL of NPB and stained with 4,6-diamidino-2-phenylindole, centrifuged at 1000xg for 10 min at 4°C, then resuspended in 1 mL of NPB. Then, nuclei were conjugated to Dynabeads™ M-280 Streptavidin (Invitrogen, Cat No./ID: 11205D) and captured on the magnetic rack. Supernatant was removed and beads were washed twice with NPB, then left resuspended in NPB on ice. The purified nuclei (25000 – 50000 nuclei) were incubated with Tagment DNA TDE1 Enzyme and Buffer mix (Illumina, Cat No./ID: 20034197) at 37°C for 30 min. DNA was purified with Wizard SV Gel and PCR clean-up system (Promega, Cat No./ID: A9282). Library was amplified using 2X KAPA HiFi HotStart (Roche, Cat No./ID: KK2601) and Nextera Primer Mix (i7+i5) for 11 cycles (98°C for 20’’, 63°C for 30’’, 72°C for 30’’; last step 72°C for 1’). Amplified libraries were purified with ProNEx Size selective Purification System (Promega, Cat No./ID: NG2001) and only libraries with fragments between 100 bp and 600 bp, peaking at 300 bp were considered. Three biological replicates were used for sequencing on HiSeq 4000 (Illumina).

### RNA-seq

For each biological replicate of RNA-seq, 25-30 seedlings grown on ½ MS plates were harvested, frozen in liquid nitrogen and ground to powder. Total RNAs were extracted using RNeasy Plant Mini Kit (Qiagen, Cat No./ID: 74904) according to the manufacturer’s instructions. Libraries generated with Illumina Stranded mRNA Prep (Illumina, Cat No./ID: 20040534) were sequenced on NovaSeq 6000 (Illumina).

## QUANTIFICATION AND STATISTICAL ANALYSIS

### ATAC-seq analyses

Reads were mapped to the A. thaliana reference genome (TAIR10) using STAR (Dobin et al., 2013). Duplicates were removed with MarkDuplicates and sorted with SortSam from Picard. Blacklisted regions as defined in (Yin et al., 2021) were removed with BEDTools intersect (Quinlan, 2014). To identify ATAC-seq peaks we employed MACS3 (Zhang et al., 2008) without shifting the reads introduced by Tn5. Consensus peaks were generated based on an overlap of at least 50% of length in all replicates. Only peaks detected in all replicates were considered and chloroplast and mitochondrial peaks were removed. ChIPseeker (Yu et al., 2015) was used for peak annotation and clusterProfiler package (Yu et al., 2012) for functional annotation of peaks. Differential accessibility analysis was done with edgeR (Chen et al., 2016). Bedgraph files created from BAM files with bamCoverage from deepTools (Ramirez et al., 2016) were scaled using –scaleFactor and converted to BigWig format with bedGraphToBigWig.

### RNA-seq analyses

Reads were trimmed with Cutadapt (Martin, 2011), filtered for ribosomal RNA with fastq_screen and further filtered for low complexity with reaper (Davis et al., 2013). Reads were aligned to the A. thaliana reference genome (TAIR10) using STAR (Dobin et al., 2013) and counted using htseq-count (Anders et al., 2015). Differential gene expression analysis was done using DESeq2 (Love et al., 2014) with the threshold of “p.adj < 0.05 and abs(log2FoldChange) > 0.6”.

### ChIP-seq re-analysis

PIF7 ChIP-seq data GSE139296 (Willige et al., 2021) in response to 4h of LRFR was re-analyzed and annotated with ChIPseeker (Yu et al., 2015). We defined PIF7 direct and shade regulated genes by overlapping PIF7 ChIP-seq and RNA-seq. H3K4me3 data from PRJNA839161 project (Calderon et al., 2022) was re-analyzed as follows, reads were aligned to the A. thaliana reference genome (TAIR10) using bowtie2 (Langmead and Salzberg, 2012), samtools (Danecek et al., 2021) was used to sort and index BAM files, and MarkDuplicates from Picard Tools to remove duplicates. Averaged bigWig files were used for plotting profiles and heatmaps with plotProfile and plotHeatmap after computeMatrix from deepTools. Published H3K9ac ChIP-seq data GSE139296 (Willige et al., 2021) was re-analyzed from averaged bigWig files as stated previously. DNA methylation (Zhou et al., 2022) profile plots for Col-0 were plotted from averaged bigWig files (GSE165001) using trackplot (Mayakonda and Westermann, 2024, Pohl and Beato, 2014).

### GO analysis

GO enrichment analysis was performed using compareCluster from the clusterProfiler package (Yu et al., 2012). Integrative Genomics Viewer was used to visualize the ATAC-seq and ChIP-seq signals (Robinson et al., 2011).

## Accession numbers for genes mentioned in this article

IAA19 – AT3G15540

YUC8 – AT4G28720

ATHB2 – AT4G16780

PIL1 - AT2G46970

PIF4 - AT2G43010

PIF5 - AT3G59060

PIF7 - AT5G61270

HFR1 - AT1G02340

HY5 - AT5G11260

HYH - AT3G17609

PHYB - AT2G18790

## SUPPLEMENTARY DATA

Supplementary Figure 1

Supplementary Figure 2

Supplementary Figure 3

Supplementary Figure 4

Supplementary Figure 5

Supplementary Figure 6

Supplementary Figure 7

Supplementary Figure 8

Supplemental table 1

Supplemental table 2

Supplemental table 3

Supplemental table 4

Supplemental table 5

## Availability of data and materials

Data generated or analyzed during this study are included in this published article and its supplementary information files. The datasets supporting the conclusions of this article are available in the Gene Expression Omnibus database, under accession no. GSE283129 (ATAC-seq, https://www.ncbi.nlm.nih.gov/geo/query/acc.cgi?acc=GSE283129) and accession no. GSE283133 (RNA-seq, https://www.ncbi.nlm.nih.gov/geo/query/acc.cgi?acc=GSE283133).

## Authors’ contributions

AB devised and performed INTACT/ATAC-seq, Western blot - SDS-Page and ChIP-qPCR. SP performed RNA-seq, Western blot - SDS-Page, ChIP-qPCR, CoP-qPCR and other experiments. RMB and MS generated INTACT lines. SP, GA, NG and RD performed bioinformatic analyses. SP and CF wrote the manuscript with contributions from all authors. All authors read and approved the final manuscript.

## Acknowledgements

We are grateful to the genome facility (GTF, https://wp.unil.ch/gtf/) for RNA-seq and ATAC-seq experiments, and to Claire Paltenghi and Jade Nicolet for their technical assistance during the screening of the transgenic lines and preliminary gene expression analyses.

