## Supplementary figures for "Transcriptional Dynamics and Chromatin Accessibility in the Regulation of Shade-Responsive Genes in Arabidopsis"

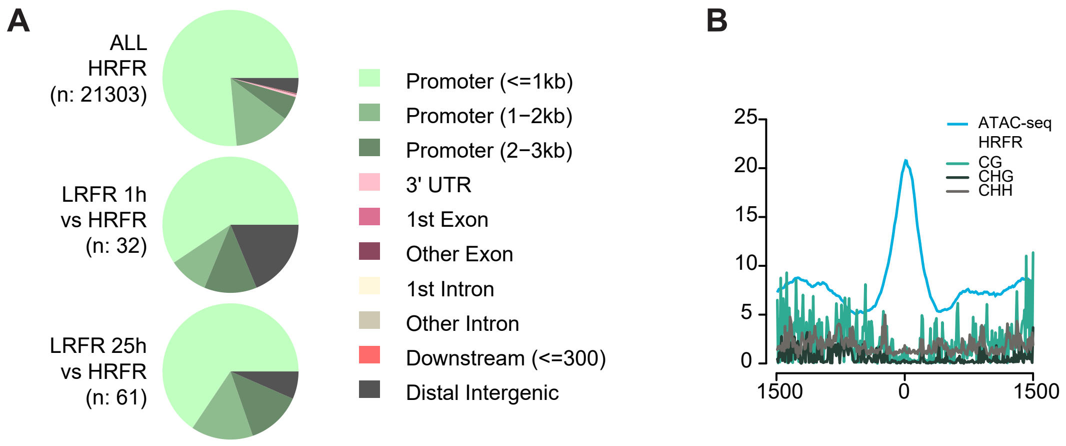


**Supplementary Figure 1. Distribution of the accessible sites in the genome.**

A. Distribution of ATAC-seq peaks across genomic regions.
B. Average ATAC-seq profile plot of 84 DARs in HRFR and average methylation profiles from (Zhou et al., 2022).

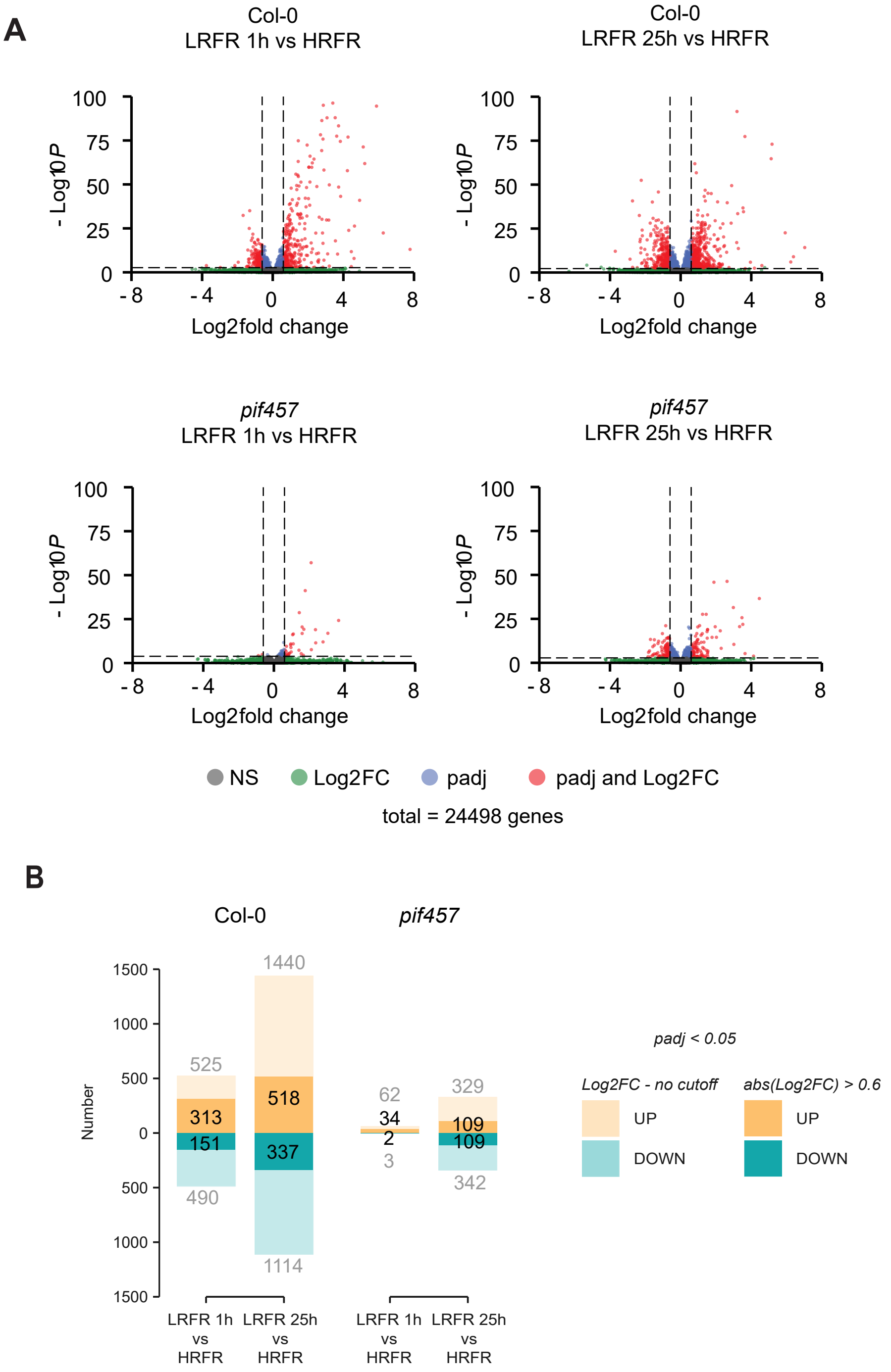


**Supplementary Figure 2. PIFs promote transcriptional response to shade**

A. Volcano plots of differentially expressed genes (DEGs) in comparisons of 1h and 25h of LRFR versus HRFR in Col-0 and *pif457* mutant. (padj < 0.05, abs(Log2FC) > 0.6).
B. Number of DEGs in comparisons of 1h and 25h of LRFR versus HRFR in Col-0 and *pif457* mutant at cutoff of padj < 0.05, Log2FC – no cutoff or abs(Log2FC) > 0.6.

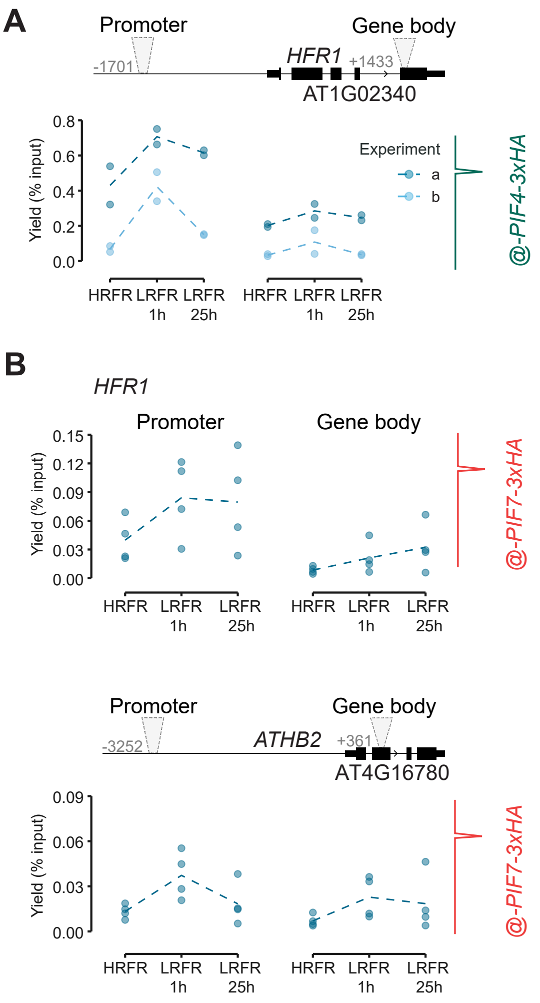


**Supplementary Figure 3. Transient increase in accumulation and stability of PIFs correlates with gene occupancy in response to LRFR.**

A. ChIP-qPCR of pPIF4:PIF4-3xHA line (in pif4-101) for *HFR1* locus (two biological replicates from two independent experiments are presented).
B. ChIP-qPCR of four biological replicates of pPIF7:PIF7-3xHA line (in *pif7-2*) for *HFR1* and *ATHB2* loci. Seedlings were grown either in HRFR for 7 days (HRFR), moved to LRFR for 1h at ZT2 of day 7 (LRFR 1h) or moved to LRFR for at ZT2 of day 6 until day 7 (LRFR 25h). Samples were collected at ZT3 on day 7.

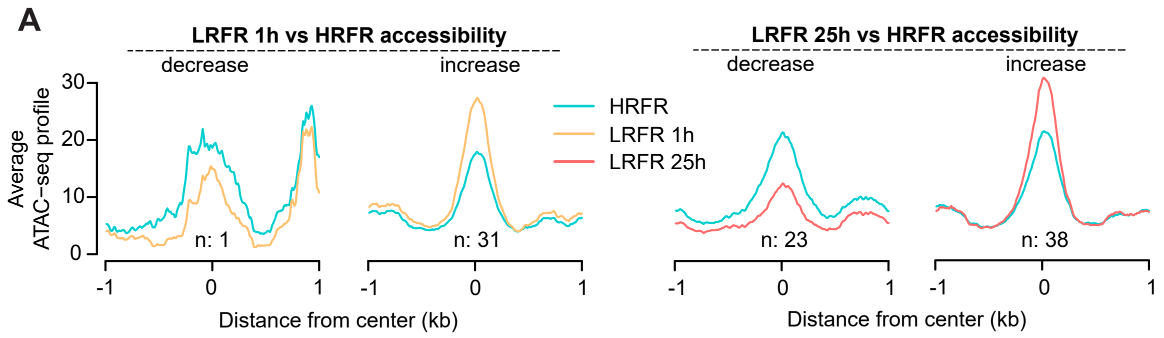


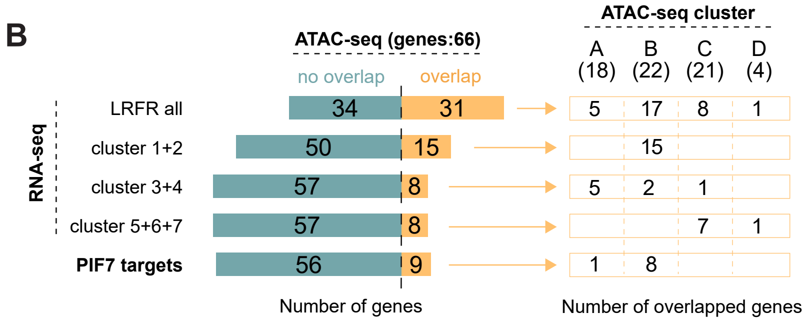


**C**

| ATAC-seq cluster | | | |
|---|---|---|---|
| A | B | C | D |
| CHIL | ACS8 | ASN1 | EXL3 |
| CLC-B | BEE1 | bZIP2 | AT1G50650 |
| GL22 | CKX5 | FLDH | AT5G01275 |
| HFR1 | CYS2 | FUM2 | AT5G12043 |
| HPL1 | GA20OX3 | HTR11 | |
| PDF2.1 | HAT2 | PLC1 | |
| SAHH2 | ATHB2 | QSOX1 | |
| TRFL1 | IAA19 | TPP | |
| TT8 | PUB25 | AT1G09157 | |
| AT1G54575 | SAUR23 | AT1G09160 | |
| AT1G65010 | ST2A | AT1G09423 | |
| AT2G01680 | AT2G18010 | AT1G26208 | |
| AT2G05995 | AT2G28060 | AT1G31540 | |
| AT2G06635 | AT2G30766 | AT1G74910 | |
| AT3G50350 | AT2G41760 | AT2G36030 | |
| AT4G13400 | AT3G16800 | AT4G16563 | |
| AT4G19500 | AT3G21330 | AT4G36850 | |
| AT5G43190 | AT3G24750 | AT5G15280 | |
| | AT4G11550 | AT5G44572 | |
| | AT4G16770 | AT5G52850 | |
| | AT4G25260 | AT5G59080 | |
| | AT5G12050 | | |

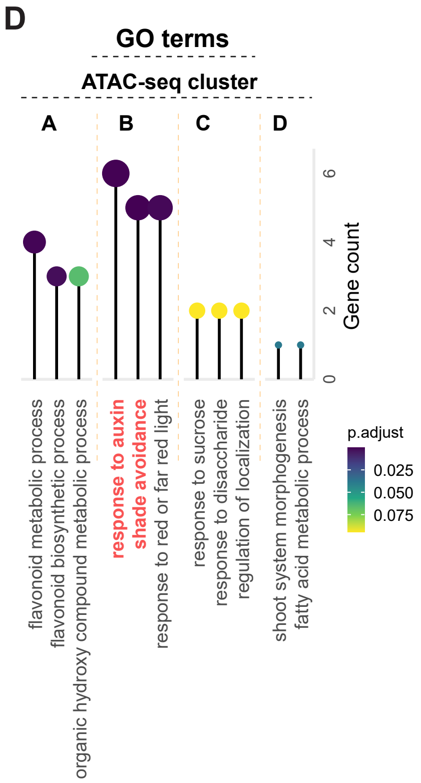


**Supplementary Figure 4. Chromatin accessibility in response to LRFR affects a moderate number of genes.**

A. Average ATAC-seq profiles of DARs in response to 1h of LRFR (left panel) and to 25h of LRFR (right panel). Distance from the center of the peak is expressed in kb.
B. Overlap of ATAC-seq genes, RNA-seq genes and PIF7 + shade regulated targets (left panel). Number of ATAC-seq genes per cluster that overlap with RNA-seq genes and PIF7 + shade regulated targets (right panel).
C. Table of ATAC-seq genes per cluster.
D. GO enrichment terms for ATAC-seq genes per cluster. The scale indicates gene counts. Adjusted P value for GO term is indicated by color scale.

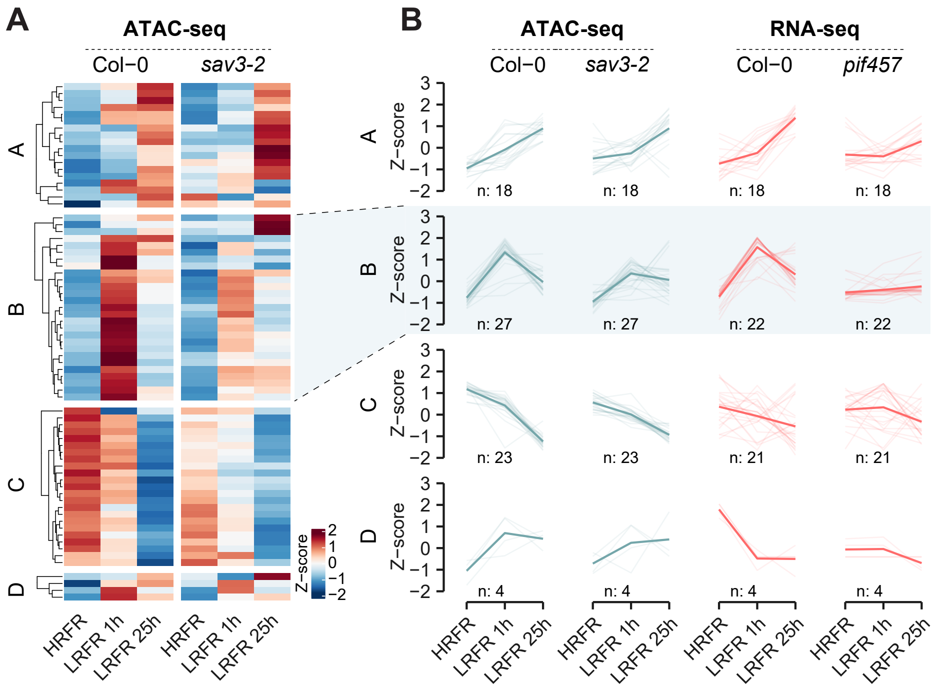


**Supplementary Figure 5. Increase in chromatin accessibility in response to LRFR in sav3-2.**

A. Heatmap of differentially accessible regions (DARs) in Col-0 and *sav3-2* in HRFR, LRFR 1h and LRFR 25h, hierarchically clustered into four distinct clusters based on ATAC-seq and RNA-seq in Col-0.
B. ATAC-seq counts of DARs in Col-0 and *sav3-2* are represented as an average z-score (in blue). The expression of genes in Col-0 and *pif457* associated with the DARs are represented as an average z-score (in red). Thick line is the average trend line. Number of DARs and genes in each cluster is displayed below the line plots.

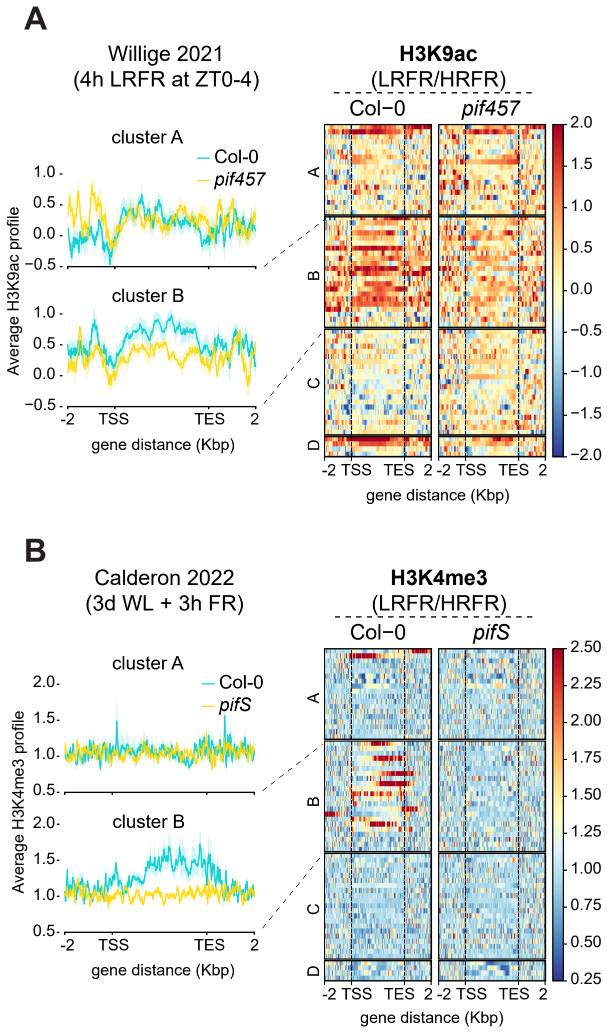


**Supplementary Figure 6. Marks of active transcription increase in PIF dependent cluster under shade.**

A. Average H3K9ac profile plot of cluster A and B genes as defined in Figure 4B (left panel) and H3K9ac heatmap across the gene bodies and +/- 2kbp regions (right panel). The data is from (Willige et al., 2021).
B. Average H3K4me3 profile plot of cluster A and B genes as defined in Figure 4B (left panel) and H3K4me3 heatmap across the gene bodies and +/- 2kbp regions (right panel). The data is from (Calderon et al., 2022).

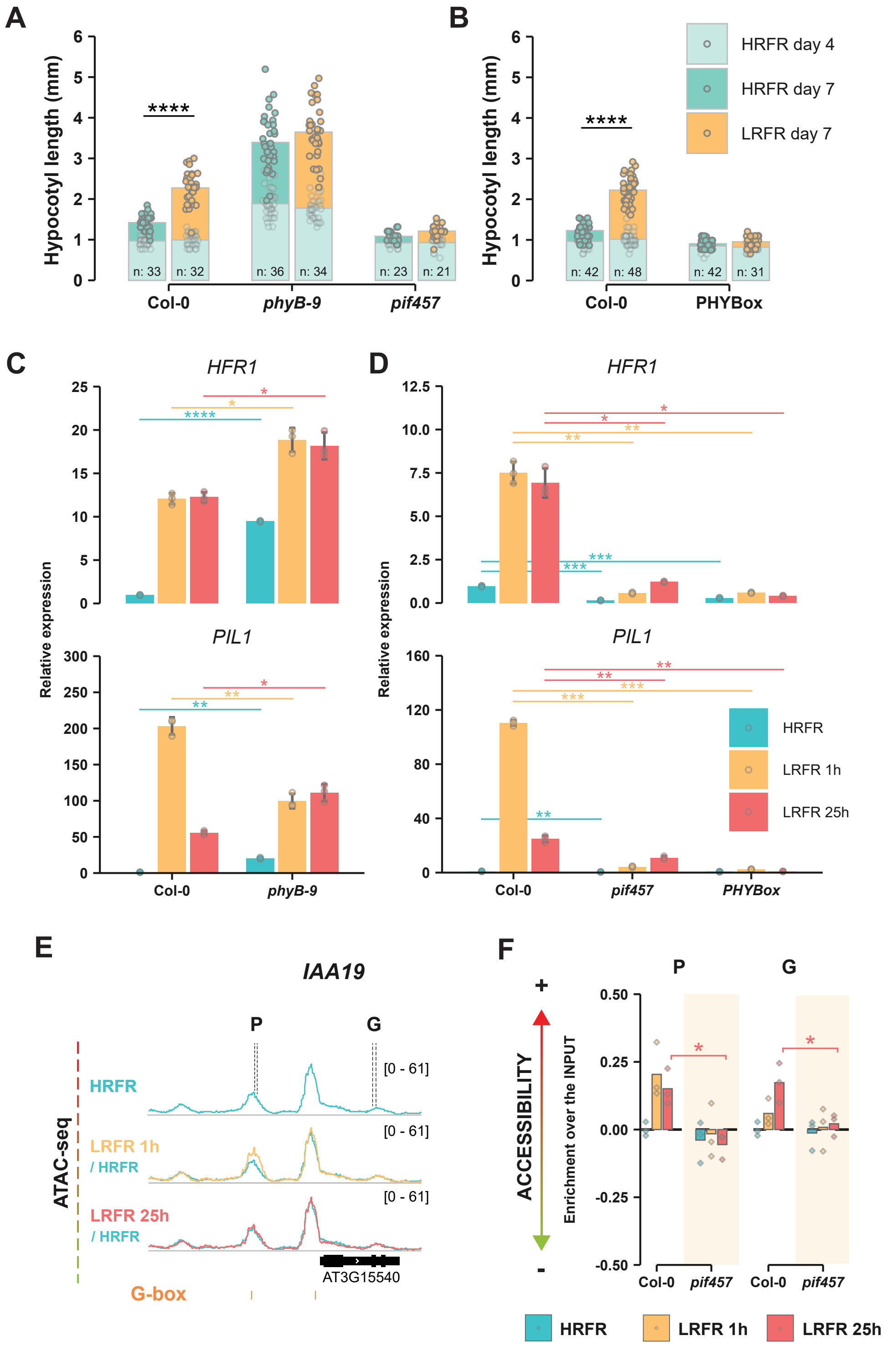


**Supplementary Figure 7. Increase in chromatin accessibility of a set of shade regulated genes is induced by PIFs in response to LRFR.**

A. Hypocotyl elongation of *phyB-9* and *pif457* mutants and
B. 35S:PHYB-GFP (PHYBox) line in response to LRFR. Seedlings were grown either in HRFR for 7 days or moved to LRFR on day 4 until day 7. Hypocotyl measurements were taken on days 4 and 7. Asterisks represent statistical significance (Students T-test, * p<0.1, ** p<0.05, ***p<0.01, ****p<0.001). s
C. Relative gene expression of PIL1 and HFR1 in phyB-9 and
D. *pif457* mutant and 35S:PHYB-GFP (PHYBox) line. Seedlings were grown either in HRFR for 7 days (HRFR), moved to LRFR for 1h at ZT2 of day 7 (LRFR 1h) or moved to LRFR for at ZT2 of day 6 until day 7 (LRFR 25h). Samples were collected at ZT3 on day 7. Asterisks represent statistical significance (Students T-test, * p<0.05, ** p<0.01, ***p<0.001, ****p<0.0001).
E. IGV view of shade regulated gene *IAA19* with changes in chromatin accessibility in response to LRFR. ATAC-seq tracks are an average of 3 biological replicates.
F. Chromatin accessibility of *IAA19* loci assayed by CoP-qPCR in Col-0 and *pif457* mutant. G-boxes are indicated in orange below the panel. Asterisks represent statistical significance (Tukey HSD, * p<0.05, ** p<0.001, ***p<0.001, ****p<0.0001).

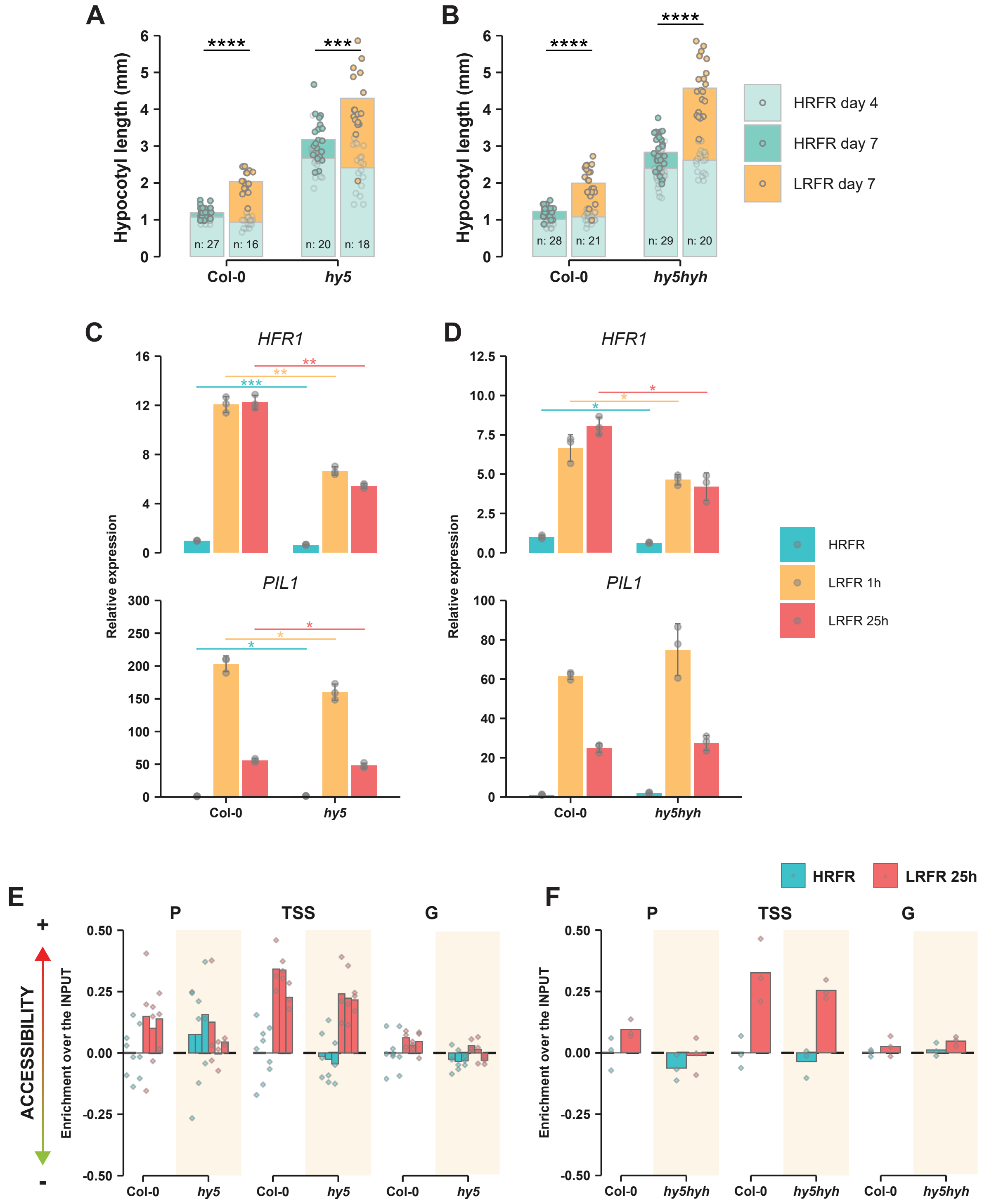


**Supplementary Figure 8. Increase in chromatin accessibility of HFR1 does not depend on HY5 or HYH.**

A. Hypocotyl elongation of Col-0 and *hy5*, and
B. *hy5hyh* mutants in response to LRFR. Seedlings were grown either in HRFR for 7 days or moved to LRFR on day 4 until day 7. Hypocotyl measurements were taken on days 4 and 7. Asterisks represent statistical significance (Students T-test, * p<0.1, ** p<0.05, ***p<0.01, ****p<0.001).
C. Relative gene expression of HFR1 and PIL1 in hy5 and
D. *hy5hyh* mutants. Seedlings were grown either in HRFR for 7 days (HRFR), moved to LRFR for 1h at ZT2 of day 7 (LRFR 1h) or moved to LRFR for at ZT2 of day 6 until day 7 (LRFR 25h). Samples were collected at ZT3 on day 7. Asterisks represent statistical significance (Students T-test, * p<0.05, ** p<0.01, ***p<0.001, ****p<0.0001).
E. Chromatin accessibility of HFR1 locus in response to 25h of LRFR assayed by CoP-qPCR in Col-0 and hy5 mutant. Three independent experiments are presented. Asterisks represent statistical significance (Tukey HSD, * p<0.05, **p<0.01, ***p<0.001, ****p<0.0001).
F. Chromatin accessibility of HFR1 locus in response to 25h of LRFR assayed by CoP-qPCR in Col-0 and *hy5hyh* double mutant. Asterisks represent statistical significance (Tukey HSD, *p<0.05, **p<0.01, ***p<0.001, ****p<0.0001).
